# Placental-expanded, mesenchymal cells improve muscle function following hip arthroplasty

**DOI:** 10.1101/297739

**Authors:** Tobias Winkler, Carsten Perka, Philipp von Roth, Alison N. Agres, Henning Plage, Bernd Preininger, Matthias Pumberger, Sven Geissler, Esther Lukasiewicz Hagai, Racheli Ofir, Lena Pinzur, Eli Eyal, Gisela Stoltenburg-Didinger, Christian Meisel, Christine Consentius, Mathias Streitz, Petra Reinke, Georg N. Duda, Hans-Dieter Volk

**Author notes:** Corresponding author: Prof. Georg N. Duda, Julius Wolff Institute and Berlin-Brandenburg Center for Regenerative Therapies, Center for Musculoskeletal Surgery, Augustenburgerplatz 1, 13353, Berlin, Germany, Charité - Universitaetsmedizin Berlin. G. N. Duda and H. D. Volk contributed equally to this study as senior authors.

## Abstract

**Background:** No regenerative approach has thus far been shown to be effective in skeletal muscle injuries, despite high frequency and associated functional deficits. We sought to address surgical trauma related muscle injuries using local intraoperative application of allogeneic placenta-derived, mesenchymal-like adherent cells (PLX-PAD), using hip arthroplasty as a standardized injury model, because of the high regenerative and immunomodulatory potency of this cell type.

**Methods:** Our pilot phase I/IIa study was prospective, randomized, double blind and placebo-controlled. Twenty patients undergoing hip arthroplasty via a direct lateral approach were injected with 3.0×10^8^ or 1.5×10^8^ PLX-PAD or a placebo into the gluteus medius muscle.

**Results:** We did not observe any relevant PLX-PAD-related adverse events at the 2-year follow-up. Improved gluteus medius strength was noted as early as week 6 in the treatment-groups. Surprisingly, until week 26 the low-dose outperformed the high-dose group and reached significantly improved strength compared to placebo, mirrored by an increase in muscle volume. Histology indicated accelerated healing after cell therapy. Biomarker studies revealed that low-dose treatment reduced the surgery-related immunological stress reaction more than high-dose. Signs of late-onset immune reactivity after high-dose treatment corresponded to reduced functional improvement.

**Conclusion:** Allogeneic PLX-PAD therapy improved strength and volume of injured skeletal muscle with a reasonable safety profile. Outcomes could be positively correlated with the modulation of early postoperative stress-related immunological reactions.

**Trial Registration:** ClinicalTrials.gov (number NCT01525667) and EudraCT (number 2011-003934-16)

**Funding:** The study was funded by the Sponsor, Pluristem Therapeutics, the Israeli innovation authority and the German Federal Ministry of Education and Research.

**Conflict of interest:** T. Winkler, C. Perka and G.N. Duda are members of a clinical advisory board of Pluristem Ltd for future indications. T. Winkler, C. Perka, G.N. Duda, P. von Roth filed a patent together with Pluristem Ltd. E. Lukasiewicz Hagai, R. Ofir, L. Pinzur and E. Eyal are current or former employees of Pluristem Ltd. T. Winkler, P. Reinke and H.-D. Volk received in the past consulting fees from Pluristem Ltd. but not for this project.

## Introduction

Skeletal muscles are of utmost importance for movement, and they also play a critical role in joint stabilization and trophic support for underlying structures. Impaired muscle functioning results in reduced strength, a reduced range of motion and joint instability, effects that are all associated with decreased quality of life, enhanced mortality, and socio-economic burden. Even so, there are no therapeutic options for regenerating injured skeletal muscles.

This problem becomes evident when the muscular abductor apparatus is injured when exposing the hip during total hip arthroplasty (THA). (*1, 2*) Rupture or destruction of the hip abductors can result in limping and predispose the joint to dislocation after THA. (*3, 4*) Any following revision substantially increases damage to the periarticular muscles. (*5*) Here, we used this highly standardized surgical procedure to assess a novel therapeutic option for muscle regeneration.

Preclinical work by us and other groups has demonstrated that the local injection of autologous mesenchymal stromal cells (MSCs) improves contraction strength after skeletal muscle injury. (*6–8*) MSCs do not differentiate into muscle cells but instead act via their secretome. (*9*) Successful regeneration results from an orchestrated interplay among stem/progenitor mobilization/activation and distinct cell-matrix interactions controlled by inflammatory processes. Therefore, the immunomodulatory effects of MSC-like cells and their secretion of trophic factors are key to tissue regeneration. (*10*) However, immune monitoring in patients that enables characterization of these effects following MSC therapy remains scarce, particularly in musculoskeletal indications.

PLacental-eXpanded mesenchymal-like adherent cells (PLX-PAD) exhibit a similar marker profile and proliferation properties as MSCs derived from other sources (*11*) but differ in their reduced ability to differentiate to mesodermal lineages. (*12*) These cells are isolated from full-term placentae and express immunomodulatory, anti-apoptotic, pro-angiogenic and anti-fibrotic (*13–15*) properties, which are key to muscle regeneration. Data from two phase I first-in-man studies in patients suffering from chronic limb ischemia demonstrated their low alloimmunogenicity, which enabled their use as a Human Leukocyte Antigen (HLA)-unmatched off-the-shelf product. (*16*) This characteristic also renders these cells ideal candidates for a clinical approach in skeletal muscle trauma.

Here, we translated our preclinical work into patient treatment using a highly standardized and frequent muscle injury. We treated damage of the gluteus medius muscle (GM) due to a direct lateral, transgluteal implantation of a total hip using a local, intramuscular injection of PLX-PAD. Patients were followed up for safety, efficacy, biomechanical muscle function, muscle morphology and the impact of the therapy on the immune system.

## Results

### Patient Demographics

Twenty-one patients were screened in the study of whom one patient was categorized as a post-randomization screening failure and not treated. We observed intra-operatively trochanteric fissures in two of the 20 patients; biomechanical analysis was not performed on these patients at visit 6 (6 weeks postoperatively). The patient demographic data are listed in Table 1. Patient flow and trial time line is shown in figure 1.

**Fig. 1:**
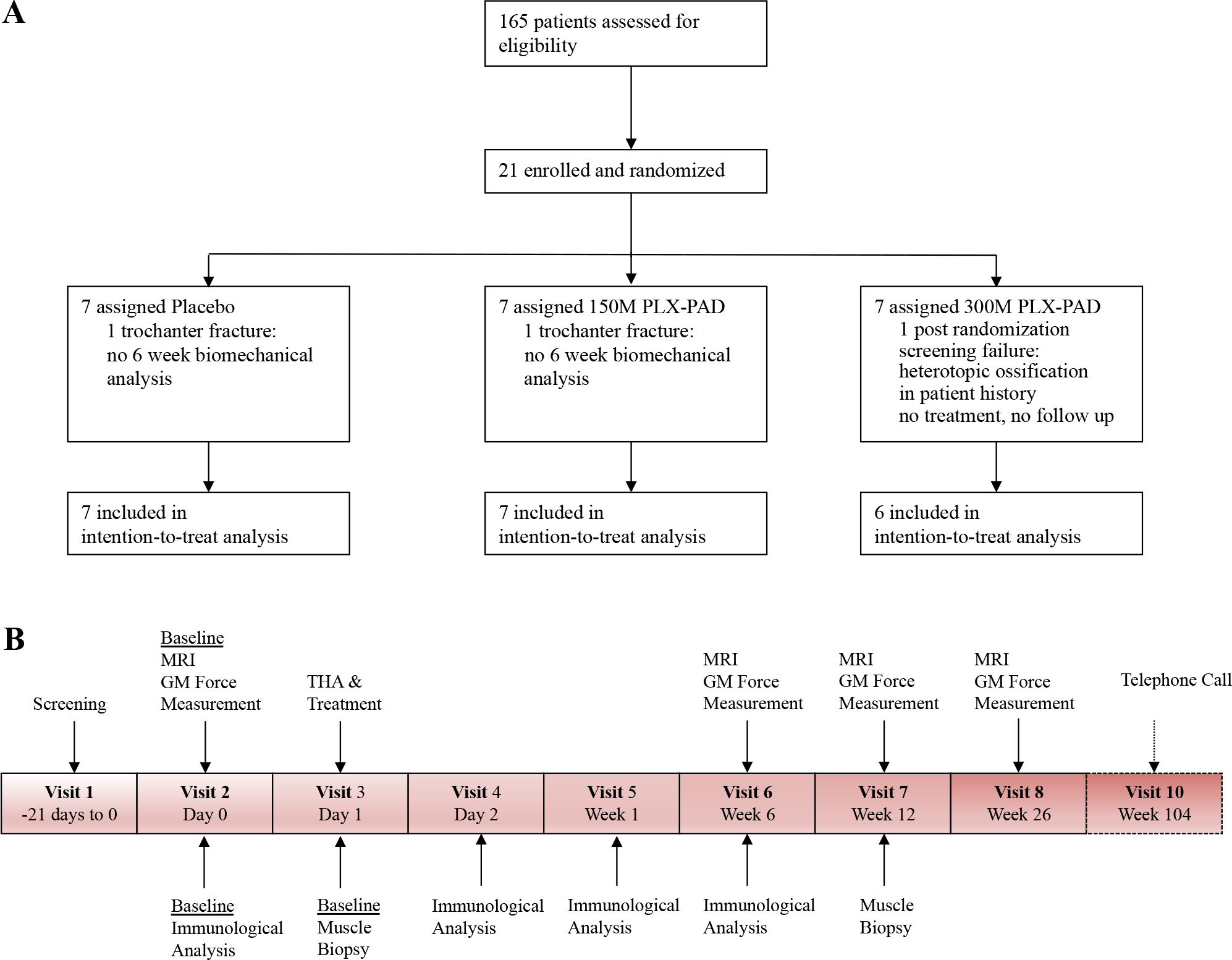
(A) Consort Diagram, (B) Timeline.

**Table 1.** Demographic characteristics of the study participants at baseline.

| Characteristics | Placebo | 150M<br>PLX-PAD | 300M<br>PLX-PAD | Total |
| --- | --- | --- | --- | --- |
|  | (n=7) | (n=7) | (n=6) | (n=20) |
| Sex, No. (%) |  |  |  |  |
| Female | 6 (85.7) | 3 (42.9) | 1 (16.7) | 10 (50.0) |
| Male | 1 (14.3) | 4 (57.1) | 5 (83.3) | 10 (50.0) |
| Age, mean (SD) | 64.3 (6.4) | 65.4 (5.9) | 64.5 (7.4) | 64.8 (6.2) |
| Ethnicity, No. (%) |  |  |  |  |
| Caucasian | 7 (100.0) | 7 (100.0) | 6 (100.0) | 20 (100.0) |
| Body Mass Index,<br>mean (SD), kg/ m<sup>2</sup> | 26.3 (2.8) | 27.5 (5.9) | 28.7 (4.2) | 27.4 (4.4) |
| Current smoker |  |  |  |  |
| No. (%) | 1 (14.3) | 3 (42.9) | 1 (16.7) | 5 (25.0) |
| years, mean (SD) | 20 | 28.7 (15.0) | 20 | 25.2 (11.6) |
| Ex smoker (years) |  |  |  |  |
| No. (%) | 2 (28.6) | 1 (14.7) | 2 (33.3) | 5 (25.0) |
| Years, mean (SD) | 26.5 (31.8) | 6 | 23.5 (16.3) | 21.2 (19.8) |

### Effect of PLX-PAD on muscle function and structure

Our primary finding was a highly significant improvement in maximal isometric contraction force in the treated abductor muscles of the low-dose group compared with the placebo group after 26 weeks (P=0.007). (Figure 2A). This improvement was accompanied by an increased GM volume (P=0.004) (Figure 3A) without evidence for an increase in intramuscular fat (Figure 3B). An enhanced contraction force was also noted on the contralateral, non-treated side in the 150M group vs. the placebo group without increasing volume (Figure 2B).

**Fig. 2.**
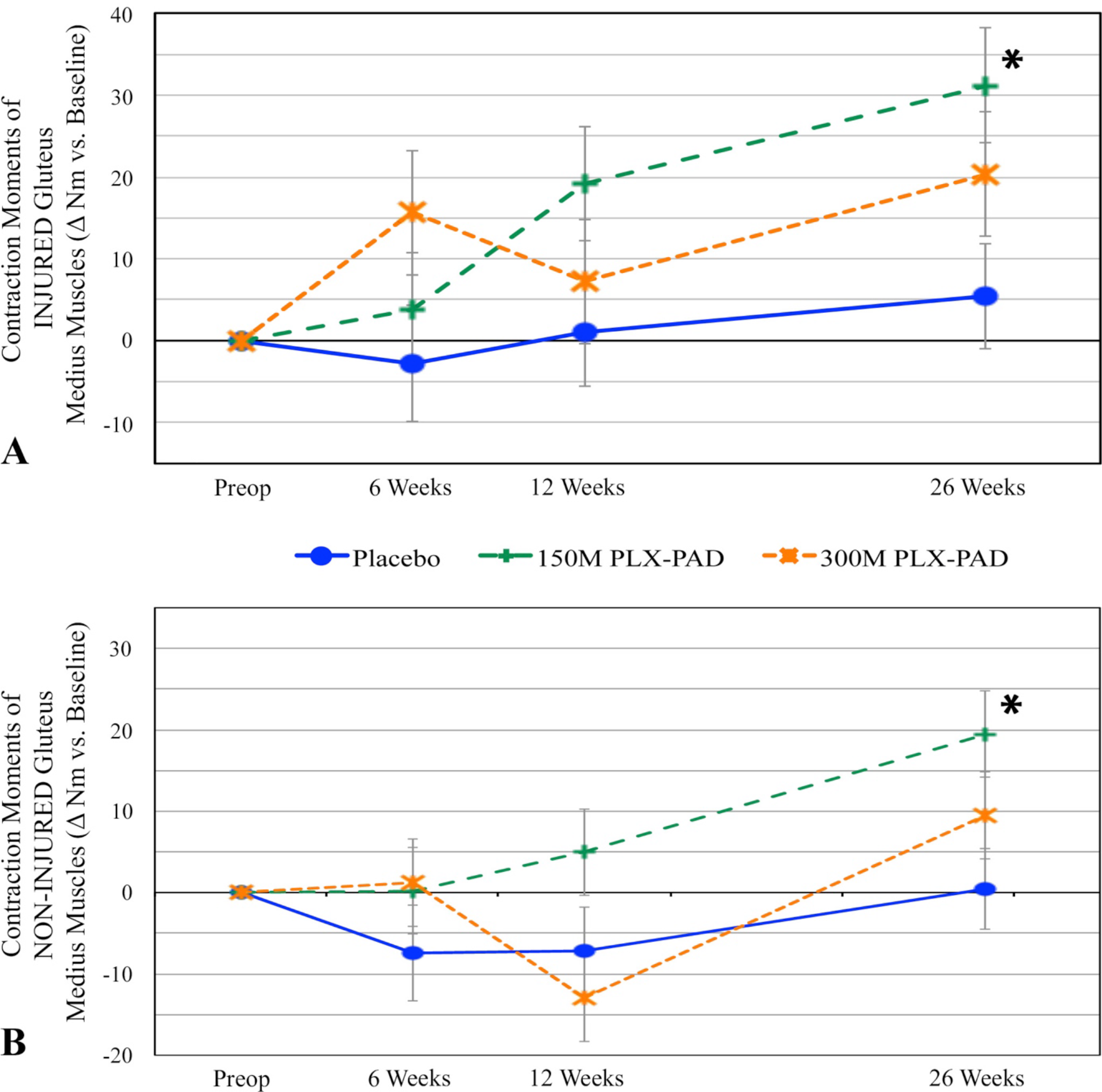
PLX-PAD treatment leads to improvement in contraction moments compared with placebo of treated and non-treated muscles. Change in mean isometric contraction moments of GM over time. (A) injured, treated side. (B) non-injured, non-treated contralateral side. Significant differences (indicated with asterisk) were found for mean isometric contraction forces in injured and uninjured muscles compared to placebo at week 26. P-value for change from baseline at week 26 150M vs. Placebo injured side: P=0.0067 (baseline adjusted; 95% CI 7.6, 43.9). P-value for change from baseline at week 26 150M vs. Placebo uninjured side: P=0.012 (baseline adjusted; 95% CI 7.1, 37.0). Repeated measures analysis of covariance, model adjusted means, modified intention to treat cohort. Preoperative baseline values of injured, treated side as mean ±SE (A) Placebo: 24.4±6.7 Nm, 150M: 27.3±5.6 Nm, 300M: 50.8±5.3 Nm. Preoperative baselines values of non-injured contralateral side (B) Placebo: 26.3±5.8 Nm, 150M: 39.5±8.4 Nm, 300M: 48.4±13.2 Nm.

The placebo group exhibited a slight increase in muscle force and volume on both sides over 6 months. In the high-dose treatment group, we observed an initial superior (treated side) or equivalent (contralateral side) muscle force compared with the low-dose and placebo group at 6 weeks, but with a decline at week 12 (Figure 2). Thereafter, the 300M group increased again but did not exceed the values of the 150M group until the 26^th^ week; this group did not reach statistical significance compared with the placebo group (Figure 2). The corresponding volume changes reflected the pattern in the force measurements of all groups, with inferior findings of the high-dose compared with the low-dose group (Figure 3A).

**Fig. 3.**
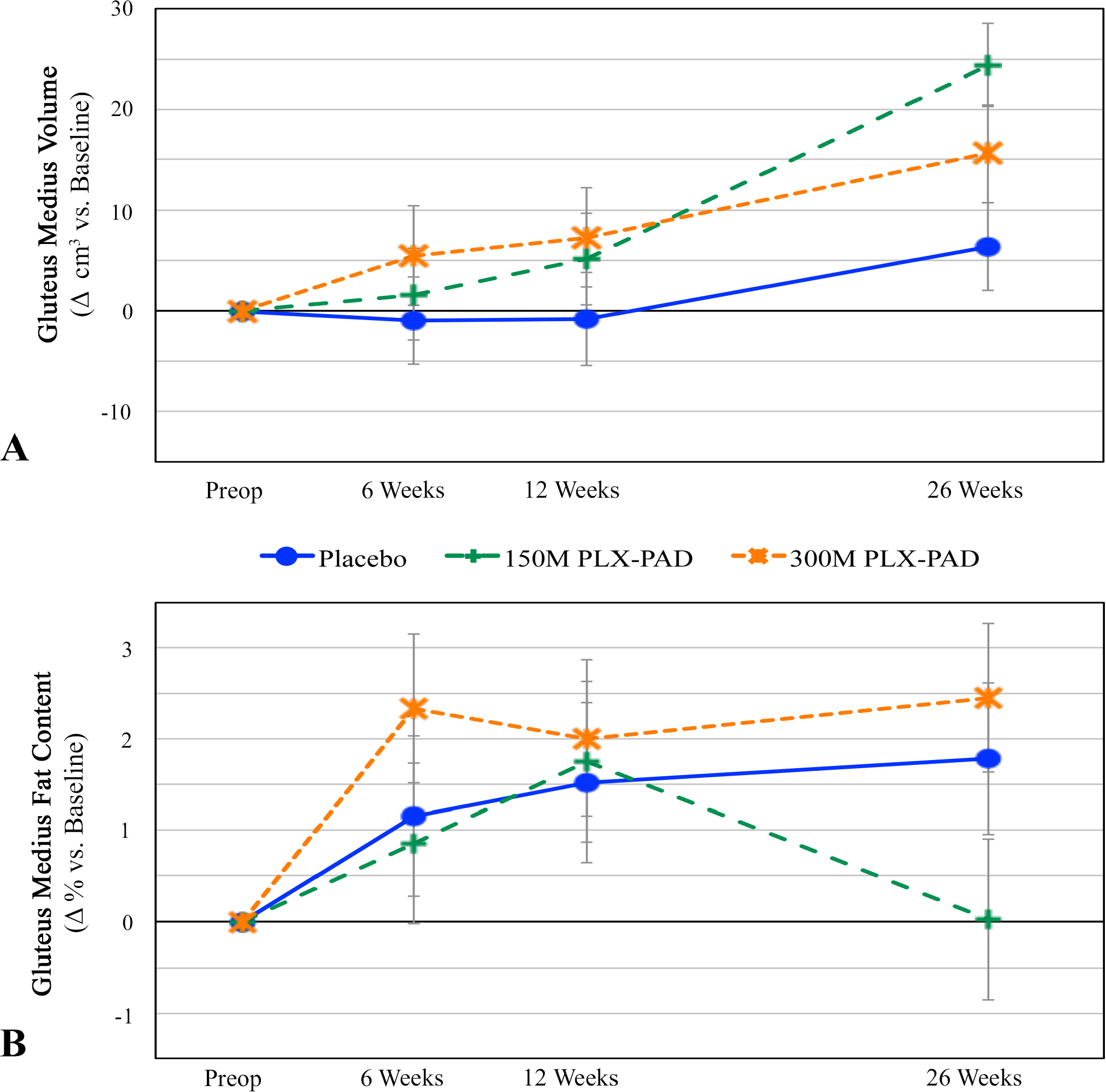
(A, B) PLX-PAD treatment increases GM volume but not fat content. Change in the macrostructure of GM over time after PLX-PAD or placebo treatment. GM volume (A) and GM fat content (B) analyses were performed via repeated MRI measurements. Significant differences (indicated with asterisk) were found for GM volume compared to placebo at week 26. P-value for change from baseline at week 26 150M vs. Placebo: P=0.004 (baseline adjusted; 95% CI 6.0, 30.0). Preoperative baseline values GM volume (A), Placebo: 211.9±15.3 cm^3^, 150M: 237.4±27.2 cm^3^, 300M: 299.5±15.0 cm^3^. Preoperative baseline values GM fat content (B), Placebo: 3.7±1.0, 150M: 6.8±2.7%, 300M: 3.3±1.3%. Δ = Change

We also analyzed whether changes in function and macrostructure were reflected on the microstructural level based on our analyses of GM needle biopsies but no significant differences were found between the groups. However, the pattern of the distribution of regenerating myofibers, as reflected in fiber diameter distribution, indicated the possibility of a faster regeneration in the cell-treated groups (Figure 4A).

**Fig. 4.**
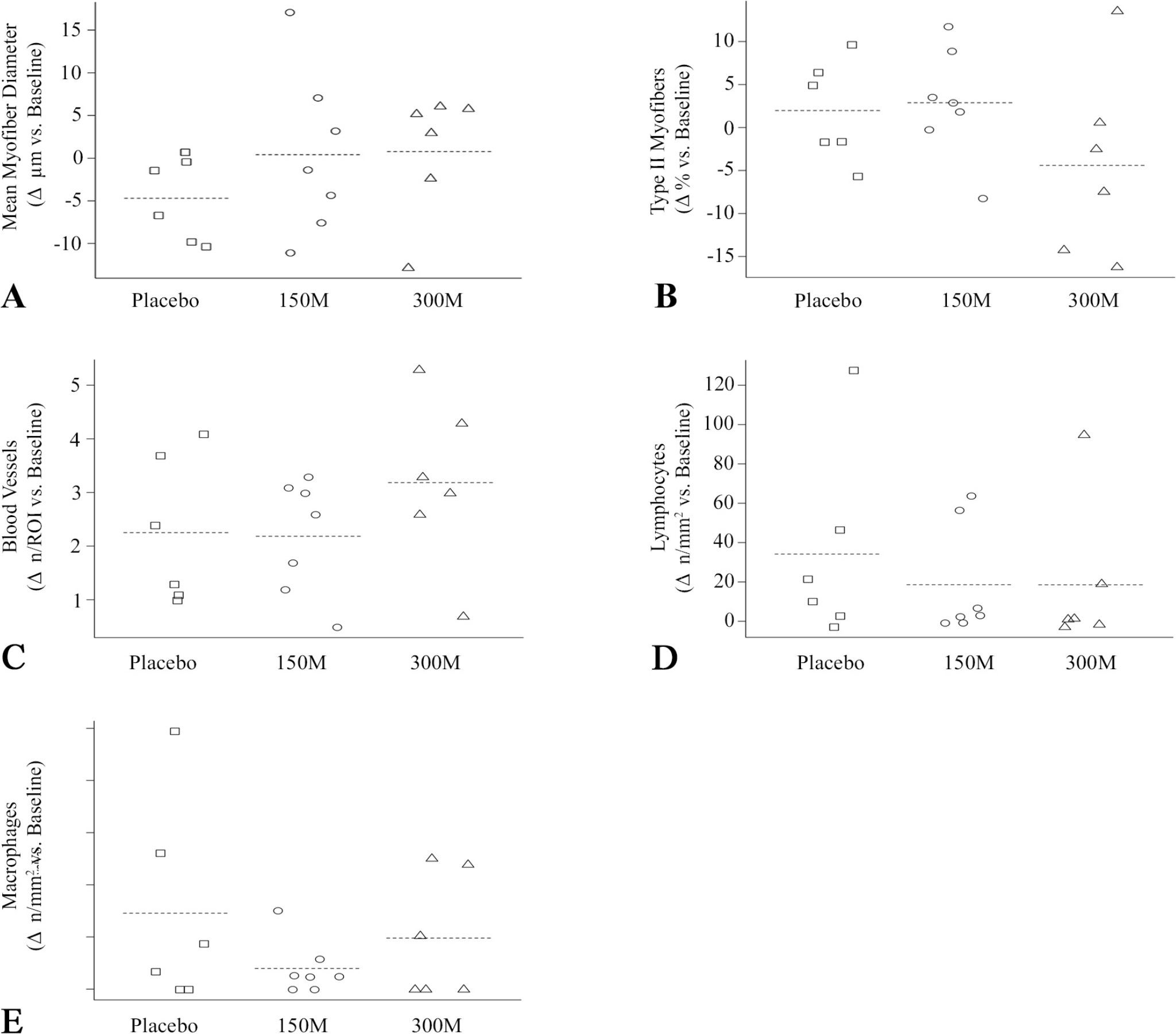
(A) Fiber distribution in PLX-PAD treated muscles indicates ongoing regeneration in placebo-treated patients. We evaluated the mean myofiber diameter (A) on muscle biopsies obtained preoperatively and 12 weeks after PLX-PAD or placebo treatment. Preoperative baseline values: Placebo: 60.5±5.1 µm, 150M: 56.9±4.6 µm, 300M: 61.7±4.1 µm. P>0.05. Δ = change. **(B, C) Myofiber type and number of blood vessels in fine needle biopsies were not changed by PLX-PAD treatment.** Preoperative baseline values myofiber type (B): Placebo: 252 ±11.5, 150M: 34.6 ±12.3, 300M: 25.1 ±16.0. Preoperative baseline values blood vessels per region of interest (ROI) (C): Placebo: 6.7 ±1.1, 150M: 5.6 ±1.1, 300M: 6.9 ±1.0. Δ = change. **(D, E) No shift of T-lymphocytes or macrophages into the gluteus medius muscle observed in biopsies.** Equal distribution of immune cells between groups within the muscle tissue. Preoperative baseline values T-lymphocytes per mm^2^ (D): Placebo: 1.3 ±1.3, 150M: 2.7 ±4.7, 300M: 5.3 ±7.5. Preoperative baseline values macrophages per mm^2^ (E): Placebo: 0.02 ±0.1, 150M: 0.04 ±0.1, 300M: 0. Δ = change. Data are given as mean±SE.

Histological analysis of fiber type change and blood vessel formation did not reveal any differences between groups in the tissue of the fine needle biopsies (Figures 4B and C).

The analysis of the local infiltration of lymphocytes and macrophages within the muscle biopsies showed an equal distribution between the groups (Figures 4D and E).

### Effect of PLX-PAD on clinical scores

We did not find any clinically relevant differences between groups in our evaluation of the Harris Hip Score (HSS) (*17*) and the Short Form-36 (SF-36) (*18*) (data shown in supplement).

### Effect of PLX-PAD on the immune system

We addressed two important questions:

i) Do PLX-PAD cells further amplify the known postoperative immune-depression related to major surgery that might pose a safety issue? Importantly, treatment did not amplify either the post-surgical immune-depression observed in the placebo group (illustrated by the strongly reduced monocytic HLA-DR expression; Figure 5A) or postoperative systemic inflammation (as in IL-6 [Figure 5B] or CrP plasma levels).
ii) Can the clear dose-dependent differences in efficacy and its kinetics be related to a differential immunomodulation by the PLX-PAD cells? Remarkably, alterations of immune cell subsets displaying the immediate postoperative stress were significantly reduced by PLX-PAD therapy, specifically in the low-dose group. This effect could be observed as a reduced relative increase in the number of CD16+ natural killer cells, activated CD19+ B-cells, CD57+8+ TEMRA cells and early IL-10. We also observed a reduced drop in the frequency of naive CD4+ T-cells and T-regulatory cell subset and reduced alterations in myeloid DC2 (CD11C+BDCA3-) cells (Figures 5C–E, supplement). Interestingly, 2 out of the 6 high-dose patients exhibited a significant increase in their TNF plasma levels within the first postoperative day.

**Fig. 5.**
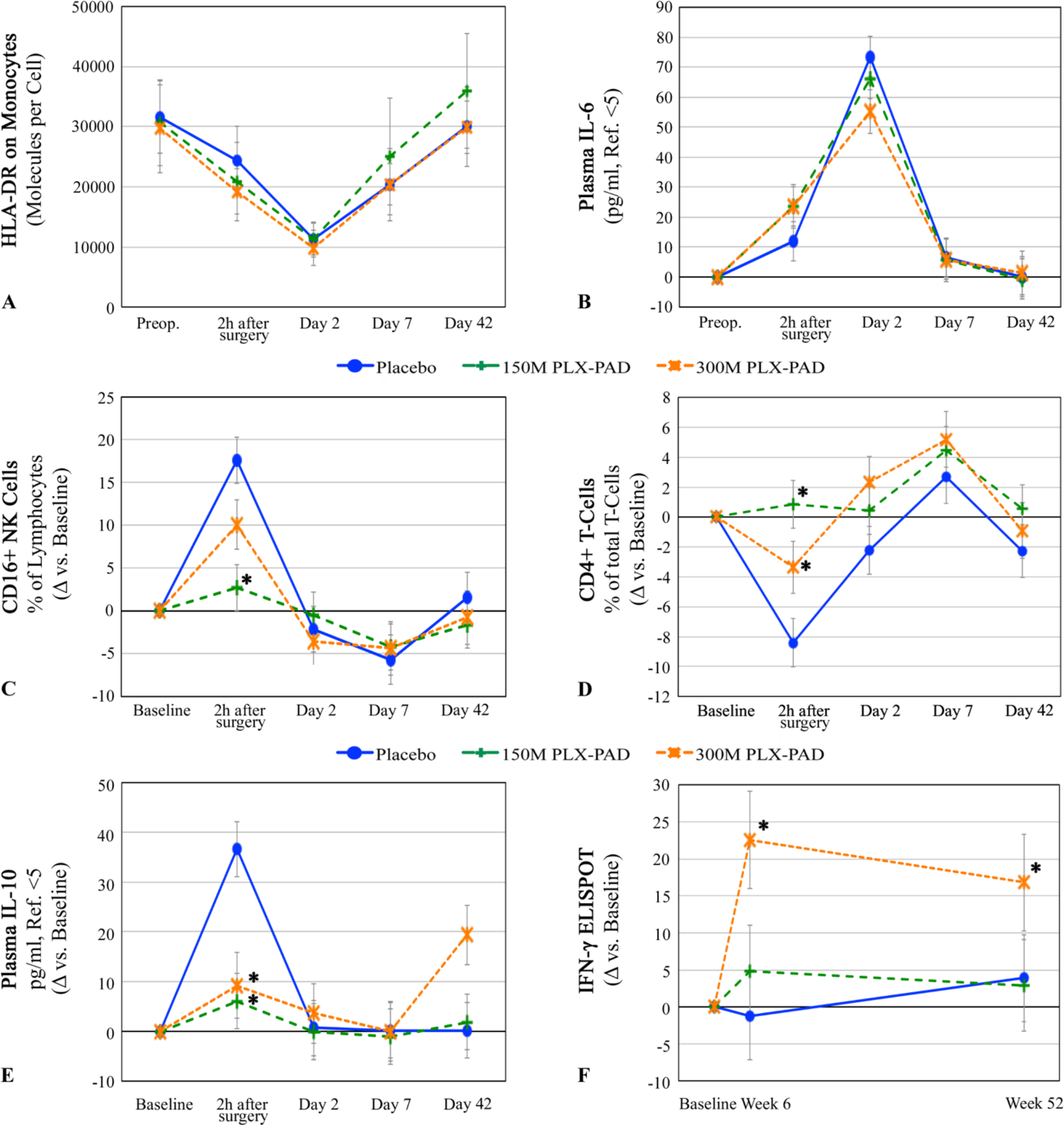
(A, B) PLX-PAD therapy does not amplify surgery-induced immune suppression. (A) Monocyte function (number of HLA-DR molecules per monocyte) shows a decrease after surgery almost below the values of immune paralysis (8000 molecules/cell), which is not altered by immunomodulation by PLX-PAD cells. (B) Plasma IL-6 levels correlate with the extent of surgical trauma and are unaffected by cell therapy. (A) and (B): total values over time. **(C, D) Prevention of surgical stress-related early changes of immune cell subset composition by PLX-PAD-therapy.** (C) 1.5×10^8^ PLX-PAD cells prevented a postoperative increase of CD16+ NK cells immediately after surgery (p<0.001) and (D) immediate postoperative decrease of CD4+ T-cells (p<0.001) High-dose group with 3.0×10^8^ PLX-PAD cells exhibited same pattern but with less pronounced effect (CD16+ NK cells: Day 1 P=0.06 vs. placebo, P=0.074 vs. 150M, CD4+ T-cells: Day 1 P=0.04 vs. placebo, P=0.08 vs. 150M). Data given as change versus baseline (mean±SE). Repeated measures analysis of covariance, model adjusted means, modified intention to treat cohort. Preoperative baseline values (C) as mean±SE: Placebo: 15.1±9.6%, 150M: 11.0±4.0%, 300M: 16.7±8.2%. Preoperative baseline values (D) (mean±SE): Placebo: 73.5±7.7%, 150M: 70.2±12.6%, 300M: 68.0±5.5%. Significant values: asterisk. **(E, F) The high-dose group displays a late rise in inflammatory parameters indicating an unspecific inflammatory reaction.** (E) PLX-PAD therapy prevented an early rise of plasma IL-10 levels, but the high-dose induced a late increase in plasma IL-10. In the low-dose group, plasma IL-10 remained at placebo levels from day 2. (F) IFN-γ ELISA demonstrated increased inflammation 6 weeks after high-dose therapy (300M vs. placebo: P=0.002), which slightly decreased until week 52 (300M vs. placebo: P=0.02). Low-dose group exhibited placebo values. Preoperative baseline values (E) as mean±SE: Placebo: 5±0 pg/ml, 150M: 6.8±4.8 pg/ml, 300M: 5.2±0.5 pg/ml. Preoperative baseline values (D) (mean±SE): Placebo: 2.6±1.9, 150M: 4.3±3.7, 300M: 1.8±1.7. Significant values: asterisk.

The histological analysis showed an equal distribution of lymphocytes and macrophages in the muscle biopsies between the groups. Therefore, it can be stated that the systemic immune changes after therapy could not be related to mere compartment shifts into the muscle tissue. However, due to the fact that not the whole muscle could be analyzed, an infiltration in other parts of the gluteus muscle cannot be fully ruled out.

Until week 6, the high-dose patient group exhibited an increased muscle force recovery (although not significant) that decreased thereafter; the low-dose group showed a slower but persistent increase and the best performance at week 26 (Figure 2). Interestingly, only high-dose patients developed significant signs of unspecific immune stimulation at week 6 corresponding to their poorer performance in terms of efficacy parameters after this time point. Figures 5E–F show rising plasma IL-10 and bystander T-cell activation for this group.

### Effect of PLX-PAD on patient safety

We did not find any safety concerns over the 2-year observation period. At week 1 after treatment, both therapy groups exhibited a transient elevation of transaminases (maximum elevations: AST, 150M: 49.3±17.7 U/l (<50); ALT, 300M: 53.7±24.3U/l (<41)), which were not detectable after 6 weeks. Treatment-emergent adverse events (TEAEs) are summarized in Table 2. Four out of the 115 TEAEs were considered to be related to study treatment (i.e., mild breath odor). Two events were considered to be serious adverse events, both of them unrelated to the study treatment.

**Table 2.**
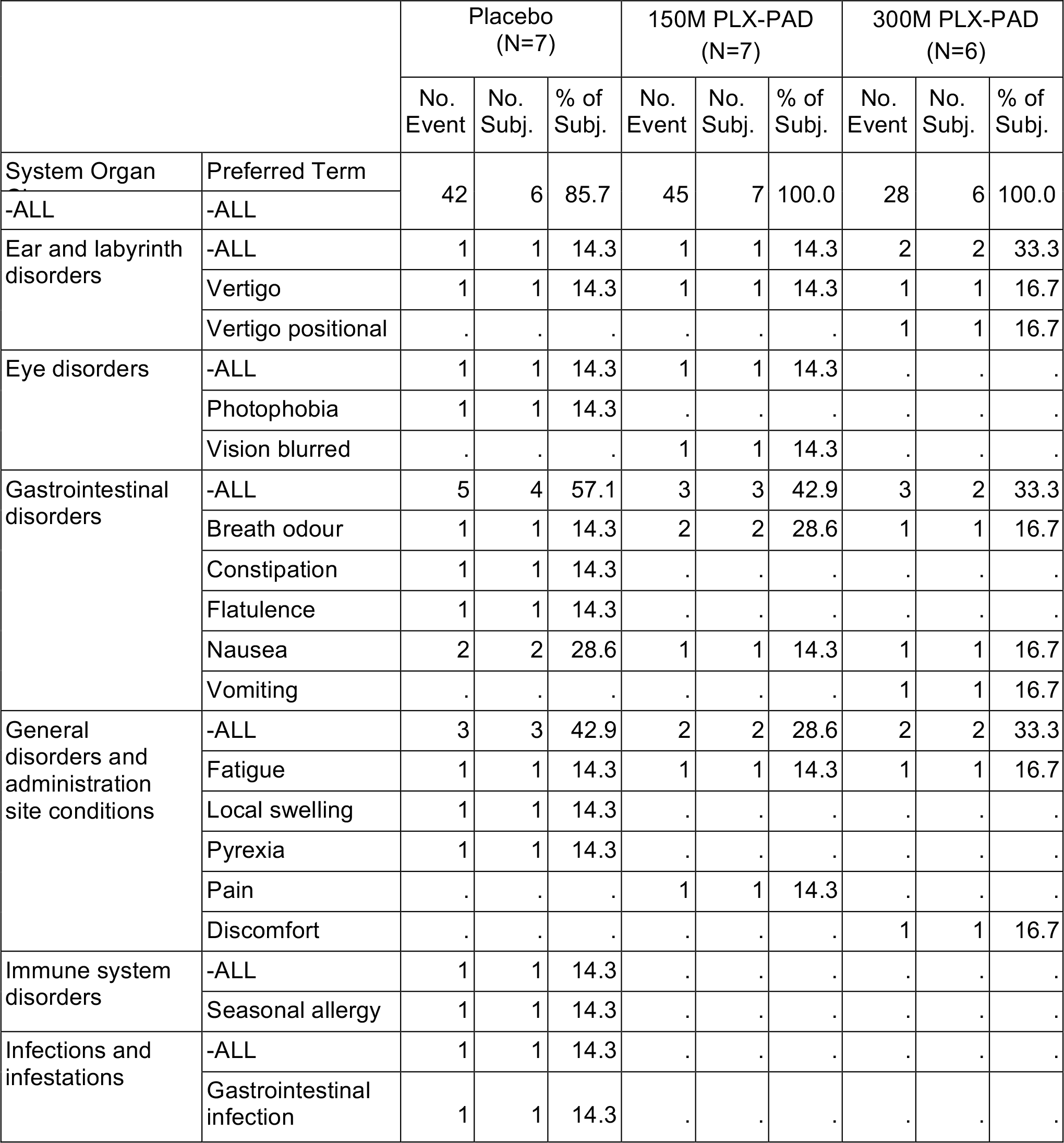
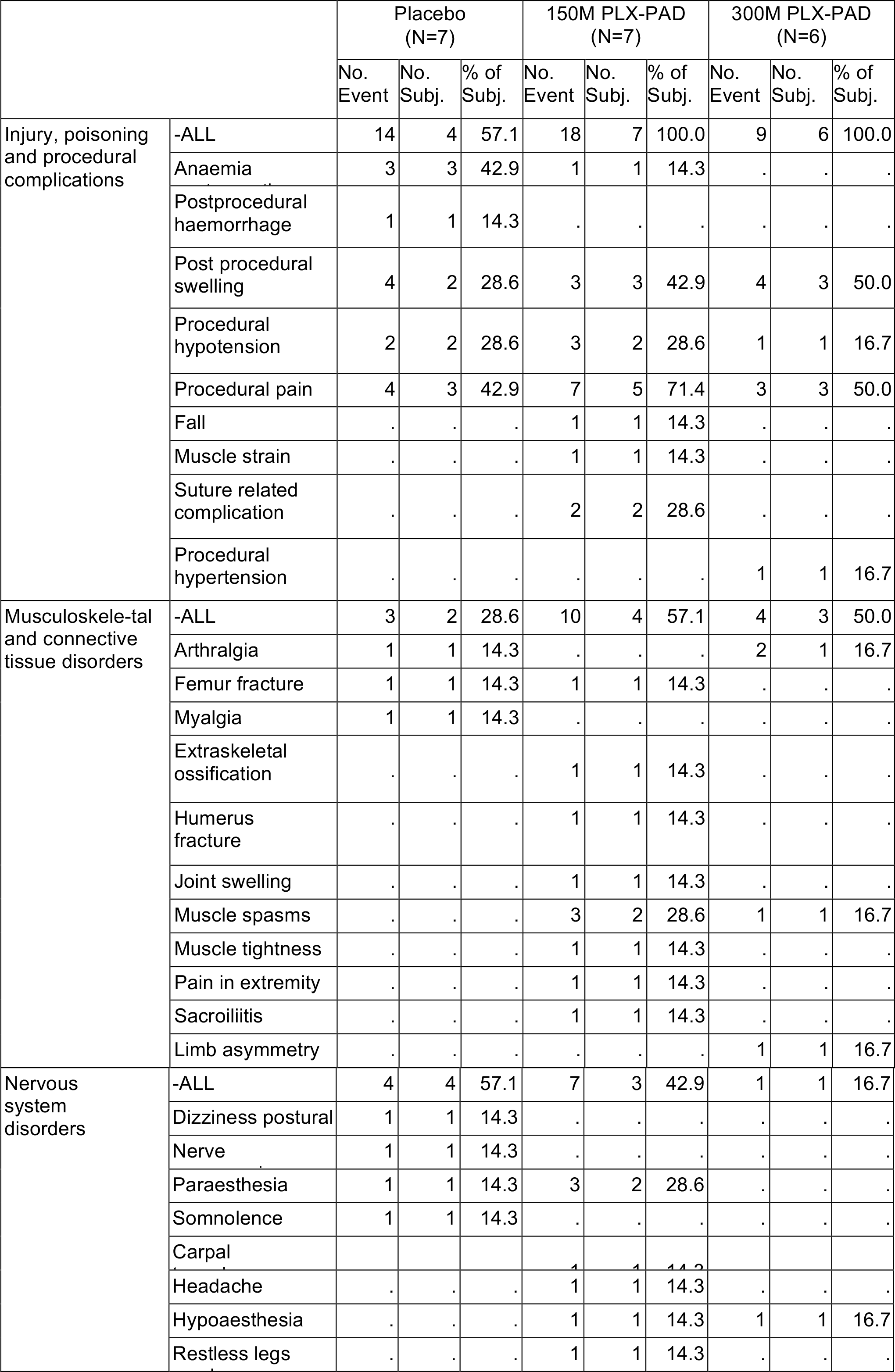
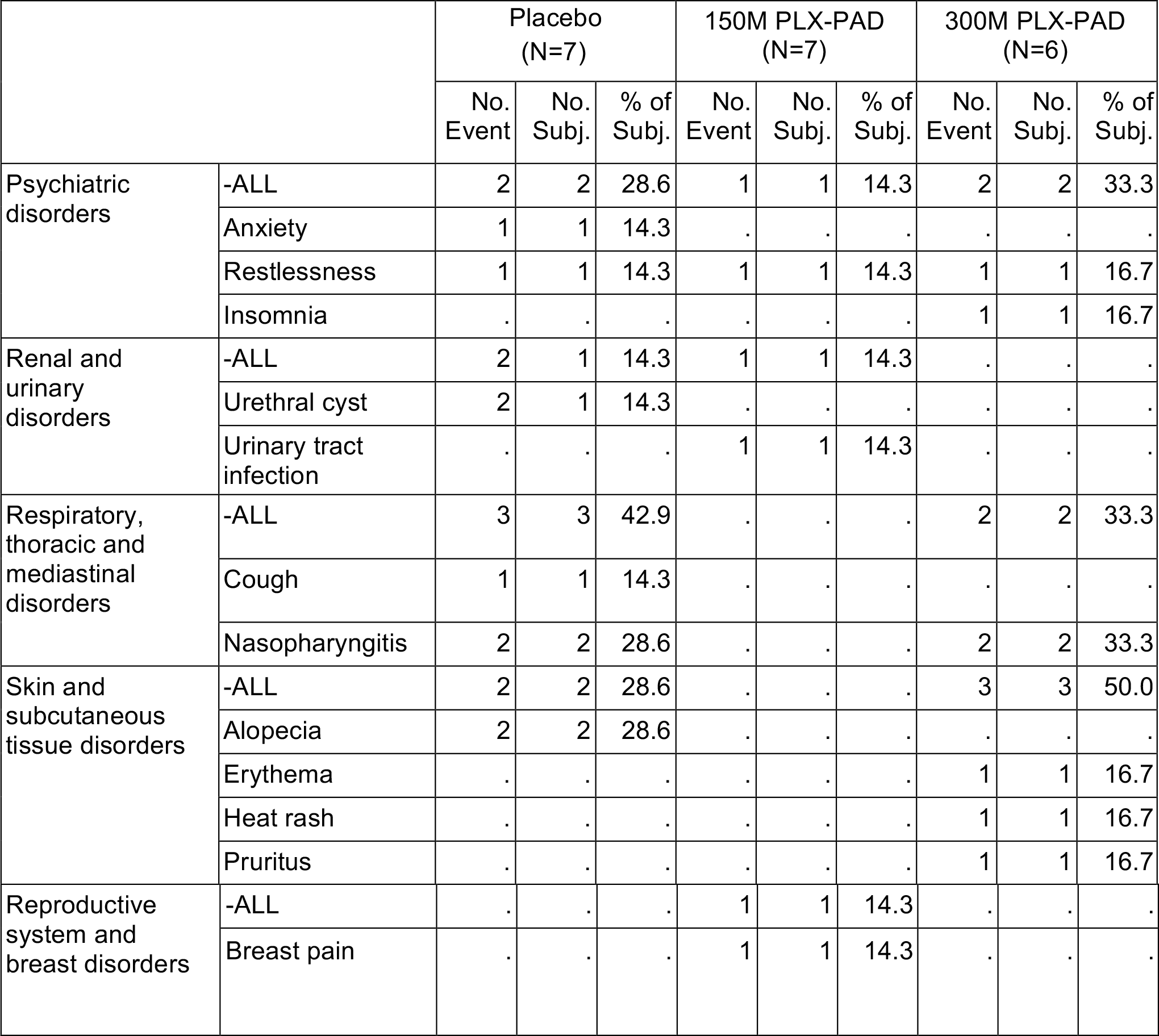
Frequency and incidence of treatment‑emergent adverse events classified by Medra System Organ Class (SOC)

Table 2 shows adverse events as classified by Medra System Organ Class (SOC) with all sub categories. All PLX-PAD treated patients and 57% of placebo treated patients had at least one of the adverse related events classified in the SOC of injury, poisoning or procedural complications. The adverse events belonging to this SOC that were observed in PLX-PAD treated patients in the clinical trial were by frequency order procedural pain, post-procedural swelling, procedural hypotension and suture related complications. In the placebo group, the AEs that were reported in the highest percent of patients were procedural pain and postoperative anemia, whereas in the PLX-PAD groups the AEs that were reported in the highest percent of patients were procedural pain and post procedural swelling. Muscle spasms/tightness occurred in a total of 4 patients treated with PLX-PAD. All these AEs were typical in the postoperative follow-up of hip arthroplasty and were categorized as not related to treatment with the PLX-PAD cells.

## Discussion

This study comprises data related to the first successful use of an allogeneic cell therapeutic approach in patients with skeletal muscle injury. We followed up patients for 2 years after treatment, and we did not observe any serious product-related side effects. The primary finding of our study was an improvement in muscle strength mirrored by an increase in muscle volume in the cell-therapy groups compared with the placebo group and an inferiority of the high-dose group versus the low-dose group. Most striking was the concordance of the results gathered from the functional assessments and micro- and macro-morphological studies with the immunological analyses.

Research of the recent past developed the concept of the immune system as a central player in regenerative processes, triggered by tissue damage. In the regeneration of skeletal muscle macrophages and T-cells coordinate proliferation and differentiation of myoblasts accompanied by a close interplay between local and systemic elements of the immune system. (10, 19, 20). In our study, histological analysis showed an equal distribution of lymphocytes and macrophages in the muscle biopsies between the groups. However, the biopsies were taken relatively late where immunological differences might be hardly detectable.

Of note was a reduced increase in muscle strength after the high-dose treatment between weeks 6 and 12 after an initial larger increase. This finding could be related to a distinct pattern in immunomodulation by low-dose treatment versus high-dose treatment as revealed by biomarker studies. Only in the high-dose group did we observe signs of late-onset immune activation in peripheral blood at week 6, which might explain the drop in efficacy in this specific dosage between weeks 6 and 12. While the peri-operative rise in IL-10 observed in the placebo group seems typical and is related to a systemic stress reaction triggering IL-10 secretion in the liver (21), the late IL-10 rise observed in the high-dose group (Figure 5E) appears to be instead a result of a counter-regulation to the ongoing inflammation by immune cells themselves. This observation is in agreement with the enhanced frequency of spontaneously IFN-gamma secreting T-cells (Figure 5F). Similar phenomena are seen post-vaccination, particularly with strong adjuvants, and are known as bystander activation. (22) The mechanisms behind bystander activation are the release of “danger” signals in association with cytokines leading to unspecific intra-tissue activation of pre-activated NK- and T-effector cells. The selected high dose (in the same volume as the low dose, meaning at higher cell concentrations) might have led to critical local conditions for the PLX-PAD cells applied and may be associated with an increased rate of dying PLX-PAD cells that would release danger signals.

Another observation was that low-dose PLX-PAD but fewer high-dose PLX-PAD almost completely abolished the surgery-associated, stress-related early changes in immune cell composition and IL-10 release. The acute stress response immediately after surgery or trauma frequently results in intestinal endotoxin translocation leading to inflammation and protein catabolism that predominantly affects skeletal muscles and results in weight loss and muscle wasting. (23) The reduction of initial protein catabolism could therefore have contributed to strength and volume advantages after the low-dose treatment over time. Paralleling the initial changes in functional parameters in the high-dose group, we observed a partial suppression of the initial stress response followed by an unspecific inflammatory reaction after the first postoperative week. The latter may be reflected in the delayed decrease in muscle strength in this high-dose group.

Our data suggest that the beneficial effects of regeneration-supporting cells are partially compromised beyond an upper dose threshold. In our preclinical studies using autologous MSCs, we observed a plateau effect in the dose-response relationship, which also indicates that a larger number of cells does not necessarily imply a better functional outcome. (*6*)

Another observation was the increase in muscle contraction force after low-dose treatment on the contralateral, non-treated side. It remains unknown, if this effect was mediated by the secretory effects of PLX-PAD cells or an improvement in neurophysiological control due to the better performance of the treated side, or a combination of both. The latter mechanism would be supported by the fact that the GM volume did not change on the non-treated side and that cross-educational effects are known to improve contralateral limb strength of the trained side by approximately 50% after unilateral training. (*24*) The effect on the contralateral side is not as pronounced as on the treated side, voting for an additional local effect of the cells.

In contrast to myoblast transplantation studies (*25–27*), our data reveal that the effect of transplanted PLX-PAD cells is not based on differentiation but rather on trophic and immunomodulatory support of the endogenous regeneration. This finding is consistent with recent studies on tendon healing demonstrating the decisive role of early trophic factor delivery to the site of injury. (*28*) Although cells were not labeled in our patients to avoid possible alterations of their biologic potential, preclinical studies had shown that transplanted PLX-PAD cells had been cleared from the host muscle within a few weeks, which highlights the importance of steering the early post-traumatic phase toward regeneration. (*29*)

Procedural pain and soft tissue swelling are typical postoperative side effects of hip arthroplasty. Analyzing their distribution in table S4, it can be seen that in the 150M group 5 patients showed procedural pain in comparison to 3 in the placebo and the high dose 300M group. We cannot exclude pain as a symptom of the early immunological effect in the 150M group, but considering that only 2 more patients stated pain in the 150M group compared to the other groups and that the 300M group was identical to placebo, we would refrain from an interpretation in this direction. Procedural swelling could be seen in only 1 more patient in the PLX-PAD groups compared to placebo, which does not allow an interpretation, in our opinion.

Main limitation of our study is the small sample size due to the pilot phase I/IIa character of the trial with primary focus on safety. Even though a small number of patients has been enrolled, the consistent results in various endpoints makes us confident about the reproducibility of these data in larger studies.

Furthermore, in depth immunological follow-up analyses revealed the clear dose-dependent effects further supporting the strength of the data despite low number of patients. Another weakness is the light sex bias with a skewed distribution between the groups, but even taking into account this bias as confounder, the support of muscle regeneration by PLX cells remains strongly significant.

In an injury, which does not comprise the whole muscle, such as the direct lateral approach used as a model in this study, the prize for the high standardization level is the compensation of contraction force within 6 weeks (first time of measurement). Patients of the placebo stayed at this functional level despite the new hip joint and an intensive rehabilitation program, which would naturally lead to an increase in force in the trained musculature. Patients treated with 150 million PLX-PAD showed an increase of contraction force and volumes of gluteus medius muscles compared to the placebo group. This increase can either be related to an improved healing of the injured muscle part (which in our opinion is the most probable based on previous work with MSCs in preclinical models), or by a hypertrophy or even hyperplasia of the non-injured muscle. Due to the restrictions of the human model a more exact differentiation was unfortunately not possible. Since we naturally could not analyze the whole muscle tissue of the patients and had to rely on fine needle biopsies of one circumscribed region of the treated muscle, the data from this analysis has to be interpreted cautiously. We did not observe an accumulation of immune cells in the treatment groups different from the placebo group and therefore did not find evidence for a compartment shift of the cells. Since we were analyzing a region that has been treated and was located directly proximal to the injury zone, we would have expected to observe an immune cell infiltration related to the treatment in this area, but, of course, an infiltration in another muscle region cannot be excluded. Likewise, we cannot exclude that another region of the gluteus medius muscle or even another periarticular muscle would show changes in vessel formation and muscle fiber type changes after treatment, but we did not observe this in the biopsies. As mentioned before we think that our immunological data may provide an explanation to why low-dose outperformed high-dose cell therapy. The central point seems to be the unspecific bystander activation of effector T cells that can migrate easily to inflamed tissue and induce an overwhelming local inflammation, which is not supportive of regeneration. Unfortunately, we were not allowed to take early biopsies, which could confirm our hypothesis derived from our immunological data. The week 12 biopsies did not reveal any differences in immune cell infiltration, which only shows that the immunological activation is resolved by endogenous regulation (e.g. IL-10) which is also reflected by the recovery of the temporary drop in muscle power increase in the high-dose group at week 12.

Promoting skeletal muscle healing is one of the remaining unsolved challenges in orthopedic and trauma surgery; there is a lack of strategies that enable complete regeneration of structure and function of muscle tissue.

In conclusion, our results demonstrate the safety of placental-expanded mesenchymal-like cells for the treatment of iatrogenic muscle injury in patients and provide preliminary results on the efficacy of this treatment. Our biomarker studies suggest that immunomodulation has a significant impact on regeneration and mediates at least partly the mode of action.

## Materials and Methods

### Study design

The study was a mono-centric, randomized, double blind and placebo-controlled phase I/IIa trial. Study site was a large academic center. We compared two dosing arms with one placebo arm (1:1:1). Randomization was based on a computorized algorithm randomizing the patients into 3 blocks of 6 patients generated by the data management of the study (CSG - Clinische Studien Gesellschaft mbH, Friedrichstraße 180, 10117 Berlin). The investigational product was shipped to the center after successful screening and randomization of a patient and thawed immediately before use. The inclusion criteria included a scheduled hip arthroplasty due to degenerative arthritis of the hip, an age of 50–75 years at the time of screening, an American Society of Anesthesiologists (ASA) score of ≤ 3 and the ability to provide informed consent. For exclusion criteria, we refer the interested reader to the supplement.

We assessed the patients for study eligibility during a 21-day screening period. The included patients were assessed for baseline parameters on day 0. On day 1, the patients underwent THA; at the end of the procedure, they received local injections of the investigational product in accordance with their allocated group. Patients in the dosing arms received 1.5×10^8^ (150M group) and 3.0×10^8^ (300M group) PLX-PAD cells, respectively.

We conducted in-clinic visits on day 2 and at weeks 1, 6, 12, 26 and 52. We assessed safety by evaluating the incidence of adverse events, assessing vital signs, physical examinations, radiological evaluations, clinical and immunological laboratory testing and electrocardiography. A telephone call at week 104 was made to inquire about newly developed malignancies.

Except for the unblinded staff members handling the treatment, all investigators, the sponsor, and any personnel involved in the subject’s assessment, monitoring, analysis, and data management (excluding the designated personnel), were blinded to the subject assignment.

The study was monitored by an unblinded and a blinded monitor. We collected the data in source documents and electronic case report forms. The data were pseudonymized and transferred to a central database. An independent data safety monitoring board conducted an overview of the study (Prof Stephan Anker, University Medical Centre Goettingen, Germany; Dr Joerg Schmidt, Klinikum Weißenfels, Germany).

### Surgery and treatment procedure

Patients underwent THA under general anesthesia via a standard lateral transgluteal approach by one surgeon (C.P.). A longitudinal skin incision was made lateral to the greater trochanter and the subcutaneous tissue dissected. A biopsy of the gluteus medius muscle was taken with a needle biopsy device (C.R. BARD Biopsie-System Magnum Core HS, Siemens, Erlangen Germany) 5 cm proximal to the greater trochanter. The iliotibial tract was opened and the gluteus medius muscle, the greater trochanter and the proximal vastus lateralis muscle exposed. Then a transgluteal approach was performed with a dissection of the periosteal flap connecting the gluteus medius and the vastus lateralis muscle followed by sharp dissection of the anterior third of the gluteus medius muscle for a distance of 5 cm proximal to the tip of the greater trochanter. The gluteus-vastus flap was then shifted anteriorly and the hip joint exposed in this fashion. Following resection of the joint capsule and the femoral head the acetabulum was reamed a non-cemented press-fit cup was implanted. The proximal femur was then exposed by transferring the leg to external rotation and adduction and a non-cemented femoral stem implanted after a further reaming procedure. The femoral head was set in place and the hip reduced. A redon drainage was set in place intra-articularly. The traumatized muscle fibers of the gluteus medius muscle were sutured with resorbable material (Vicryl, Ethicon, Somerville, USA) in single node technique. Following this, the investigational product was thawed and administered into the muscle in 10 injections of 1.5 ml around the cut, in the distance of 1 cm from the cut prior to wound closure. With the used volume and the separation in 10 single injections we were in every case able to yield depots in the muscle without reflux of the treatment fluid.

The patients were mobilized from the first postoperative day with full weight bearing and received a standardized rehabilitation program during their clinical stay and upon their discharge on postoperative day 8.

### Efficacy endpoints

The primary efficacy endpoint was the function of the GM muscle quantified by the change in maximum voluntary isometric contraction force (MVIC) from baseline following treatment. Additional efficacy endpoints included the change in total GM volume and fat content, GM fiber type and diameter, pain, Harris Hip Score (HHS) and quality of life assessed using the Short Form-36 (SF-36).

We performed maximal isometric force measurements of the GM bilaterally on a dynamometer preoperatively and at weeks 6, 12 and 26 postoperatively.

We calculated GM volume and fat content from MRI measurements at identical time points. Fine-needle biopsies were obtained from the GM 5 cm proximal to the major trochanter of the treated hip during surgery and 12 weeks thereafter.

### Biomechanical analysis

Isometric hip abductor strength of each limb was assessed using a dynamometer operating at 1000 Hz (Biodex System 3, Biodex Medical Systems, Shirley, NY), on which maximal voluntary isometric contractions (MVICs) were performed to determine the maximal voluntary torque. Prior to measurements, patients were warmed up by walking for at least 15 minutes. The test procedure was explained in detail to each patient, with special attention paid to the positioning and execution of the contraction by an experienced physiotherapist. A custom-made support was used to support the non-tested limb during each test, to maintain a proper position with the body upright. All patients were tested in a standing position facing the dynamometer with the legs shoulder width apart, with the hip rotation axis aligned with the dynamometer axis. For familiarization purposes, patients performed a submaximal contraction prior to data collection. Each subject performed an MVIC for 15 seconds with at least 60s rest between each MVIC.

Similar techniques to assess hip abduction strength by measuring the torque moment using a dynamometer in a standing position have been used in sports science studies, in both isokinetic (*30*) and maximal fatigue (*31*) tests. This test has previously been shown to be reliable in maximal isometric tests, with excellent test-retest reliability (ICC = 0.917 between separate sessions) for 15-second maximal hip abduction efforts. (*31*)

### MRI analysis

The evaluation of volume and fatty atrophy of the gluteus muscles was performed on axial T1- weighted MR Images by two blinded investigators on a PACS workstation (Osirix (Geneva, Switzerland) and Image Software Photoshop (San Jose, USA)).

For the volume analysis, each muscle was segmented in the transversal plane creating polygons with a slice thickness of 5mm. The total muscle volume was calculated from the sum of volumes of all polygons received from a muscle. (Figure 3A).

The analysis for fatty infiltration was performed using a technique described previously (*5, 32*). Briefly, three slices 30mm proximal to the trochanter major were selected in each patient. On the basis of regions of interests in the subcutaneous fatty tissue and the ipsilateral iliac muscle the mean intensity of gray levels were defined defining fat or muscle tissue. The percentage of fatty infiltration in the gluteus muscle was calculated by using the ratio of pixels of fat-value-and muscle-value-pixels in the determined slices. (Figure 3B)

### Histological analysis

All muscle samples were snap frozen, embedded in Tissue-Tek^®^ O.C.T.TM compound and cut into 10 µm cross-sections (Microm HM 60, MICROM International, Walldorf, Germany). The frozen sections were thawed and stained hematoxylin and eosin (Merck, Germany) or immunhistochemistry (IHS) was performed. Briefly, for the IHS, sections were 30-min air-dried, fixed and washed in PBS. Before the primary antibodies were used, all sections were incubated with blocking solution (horse or goat serum Biozol, Eching, Germany) for 30 min. IHC stains were performed with biotinylated antibodies against fast myosin heavy chain (fast-MHC, clone My 32, #M4276, Sigma-Aldrich, St. Louis, USA), Factor VIII (#CP 039B, Biocare Medical, Concorde, USA), αβ-t-cell-receptor (#SM1230PS, Acris Antibodies GmbH, Herford, Germany) and CD68 (#SM1718T, Acris Antibodies GmbH, Herford, Germany). After a 60-minute incubation with the primary antibody at room temperature, sections were washed twice with PBS for 5 minutes. Afterwards the secondary antibody was applied (anti-mouse or anti-rabbit, VectorLaboratories Inc, Burlingame, USA) for 60 minutes and the sections were washed again. Finally, the avidin-biotin-complex (# AK 5000, Alkaline Phosphatase Standard Kit, Vector Laboratories, Burlingame, USA) and Vector®Red Substrat Kit (Vector Laboratories, Burlingame, USA) for signal amplification were applied and nuclei were counterstained acoording to the Mayers Hemalum method.

All images were investigated with a light microscope (Leica Microsystems, Germany) equipped with a digital camera (AxioCam MRc, Carl Zeiss MicroImaging, Germany) in a hundredfold magnification. Subsequently sections were inverted to an entire image using the Axiovision software (Zeiss, Göttingen, Germany). For the quantification of the histomorphometric parameters two blinded investigators analyzed the samples manually by using regions of interests (ROIs) generated randomly or the total sections as described below using the software ImageJ (Maryland, USA). For the calculation of the mean myofiber diameter the shortest diameter of 400 fibers in ROIs on H&E sections were measured and the mean calculated for each patient (Figure 4A). Type II and type I myofibers were counted on total sections stained for fast-MHC and the percentage of type II of total fibers calculated (Figure 4B). Blood vessels were counted in 10 random ROIs of 500 x 500 pixels and given as all FVIII positive vessels per ROI (Figure 4C). T-lymphocytes were counted on total sections as αβ-t-cell-receptor positive cells and given as cells per mm^2^ of muscle tissue. Lymphocyte numbers in clusters as depicted in Figure 4D left were calculated via the ratio of area of the cluster divided by the mean diameter of a lymphocyte nucleus, previously determined by measuring 50 random nuclei on the sections as 10.2 µm. Macrophages were counted as CD68 positive cells on total sections and given as cells per mm^2^ of muscle tissue (Figure 4E).

### Immunological analysis

We monitored the patients in terms of their immune cell subset composition, cytokine and endothelial activation marker plasma levels, and ex vivo monocyte and T cell function. We obtained blood samples preoperatively, 2 hours after surgery and at days 2, 7 and 42 postoperatively. For additional details about the procedure, we refer the interested reader to the supplementary information.

### Test compound

PLX-PAD is an allogeneic ex-vivo placental expanded adherent stromal cell product. The mesenchymal-like stromal cells, termed adherent stromal cells (ASCs) have been derived from the full term human placenta following a caesarean section and expanded using plastic adherence on tissue culture dishes followed by 3-dimensional (3D) growth on carriers in a bioreactor. Seeding the cells on fibra-cel disks and placing them in the bioreactor provides a 3D-structure microenvironment that enables controlled large-scale growth of these cells. PLX-PAD cells obtained from Pluristem Ltd. are stable adhesive cells that can be expanded in vitro without the loss of phenotype and without showing signs of karyotypic changes. PLX-PAD are spindle in shape with a flat, polygonal morphology, and 15-19 µm in diameter.

PLX-PAD cells were further characterized in our institute by in-depth surface marker analysis. For this purpose, we applied the “Human Cell Surface Marker Screening (PE)” Kit (Biolegend) using directly labeled antibodies for detecting surface markers. We compared several batches of PLX-PAD cells with a bone-marrow derived MSC line. Table 3 summarizes the data from the CD screen. PLX-PAD cells showed the consensus expression profile of MSC, such as CD 73+ 90+ 105+ CD45-31-34-. In line with the unique properties of PLX-PAD cells compared to conventional MSC, however, they show a much broader expression profile. In summary 43 of 243 surface markers screened were expressed by PLX-PAD cells revealing a unique profile. Remarkably, there is a broader expression of various adhesion molecules (e.g. CD49 family, CD144), inhibitors of complement activation (e.g. CD46, CD55, CD59) and T-cell function (e.g. PD-L), molecules involved in signal transduction (e.g. CD140b, CD150), and molecules interacting with metalloproteinases (e.g. CD 10) but missing some markers expressed on bone marrow derived MSC which are involved in cell activation (e.g. CD109, CD112). This expression profile is in line with the proposed anti-inflammatory, immunomodulatory, cell-interaction, tissue homeostasis and angiogenesis influencing properties of PLX-PAD. Importantly, PLX-PAD cells from different preparations or even from different donors expressed an almost identical marker profile underlying the robustness of the manufacturing process (not shown).

**Table 3.**
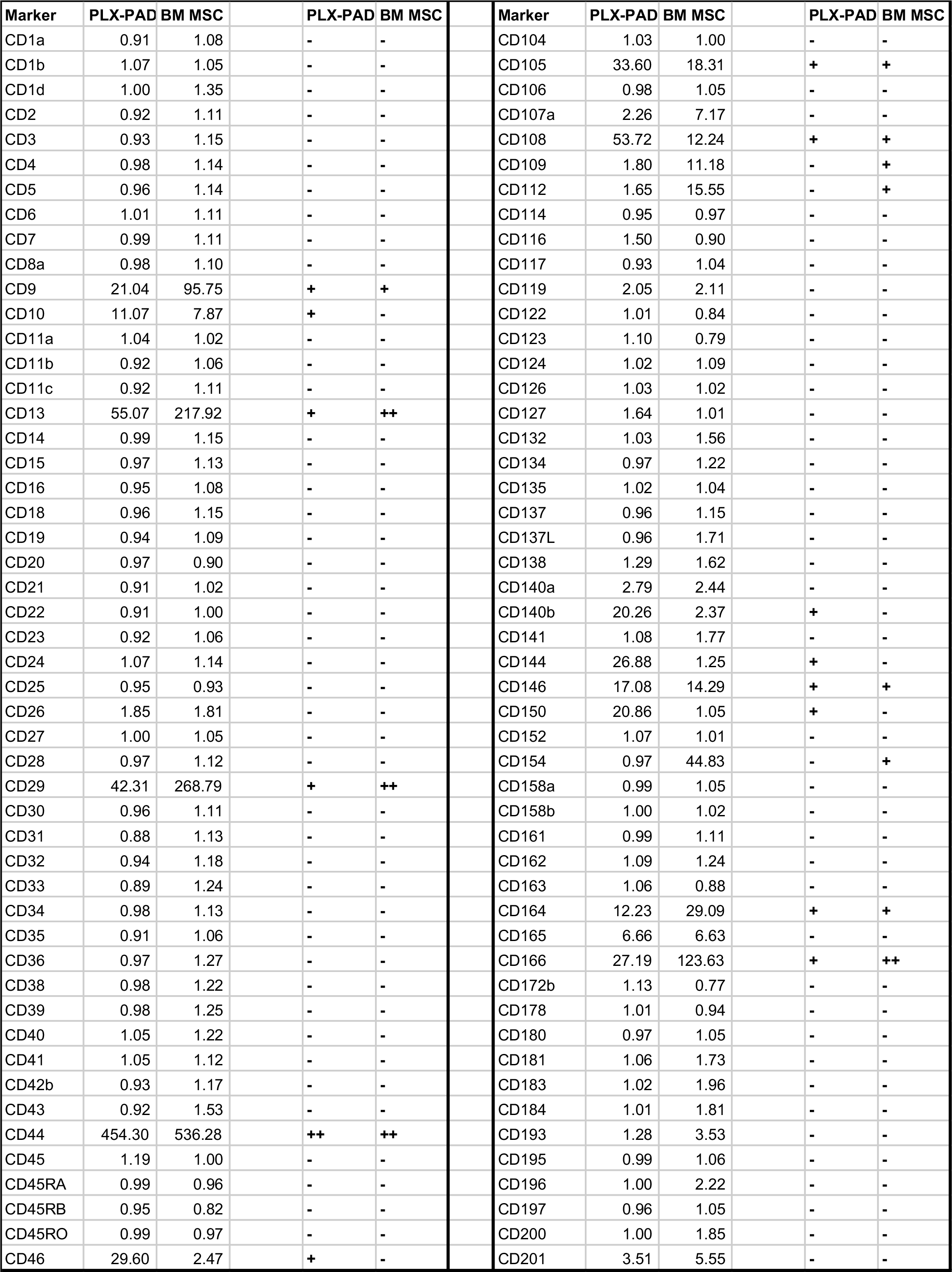
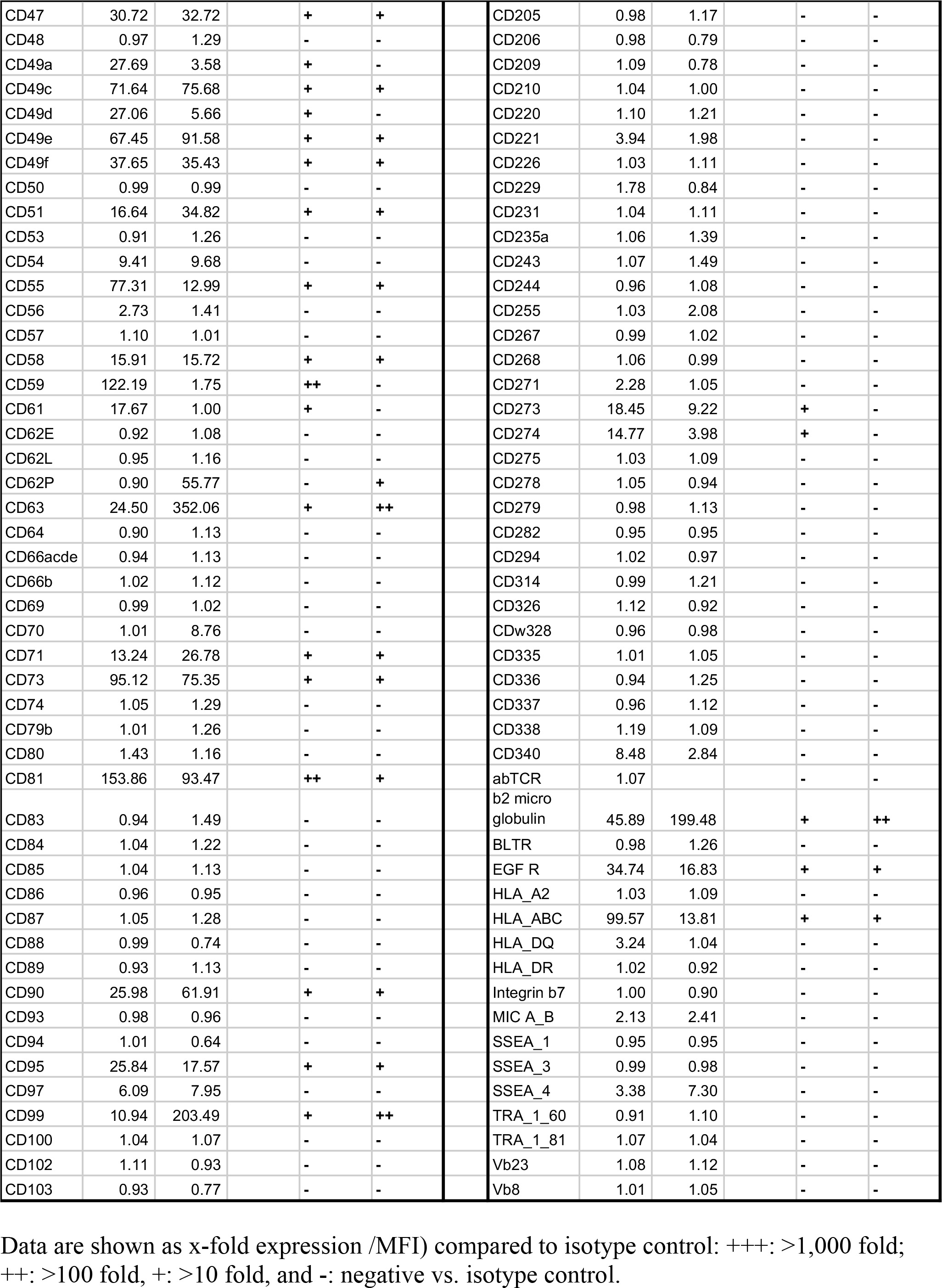
Characteristics of PLX PAD cells.

| Marker | PLX-PAD | BM MSC |  | PLX-PAD | BM MSC |  | Marker | PLX-PAD | BM MSC |  | PLX-PAD | BM MSC |
| --- | --- | --- | --- | --- | --- | --- | --- | --- | --- | --- | --- | --- |
| CD1a | 0.91 | 1.08 |  | - | - |  | CD104 | 1.03 | 1.00 |  | - | - |
| CD1b | 1.07 | 1.05 |  | - | - |  | CD105 | 33.60 | 18.31 |  | + | + |
| CD1d | 1.00 | 1.35 |  | - | - |  | CD106 | 0.98 | 1.05 |  | - | - |
| CD2 | 0.92 | 1.11 |  | - | - |  | CD107a | 2.26 | 7.17 |  | - | - |
| CD3 | 0.93 | 1.15 |  | - | - |  | CD108 | 53.72 | 12.24 |  | + | + |
| CD4 | 0.98 | 1.14 |  | - | - |  | CD109 | 1.80 | 11.18 |  | - | + |
| CD5 | 0.96 | 1.14 |  | - | - |  | CD112 | 1.65 | 15.55 |  | - | + |
| CD6 | 1.01 | 1.11 |  | - | - |  | CD114 | 0.95 | 0.97 |  | - | - |
| CD7 | 0.99 | 1.11 |  | - | - |  | CD116 | 1.50 | 0.90 |  | - | - |
| CD8a | 0.98 | 1.10 |  | - | - |  | CD117 | 0.93 | 1.04 |  | - | - |
| CD9 | 21.04 | 95.75 |  | + | + |  | CD119 | 2.05 | 2.11 |  | - | - |
| CD10 | 11.07 | 7.87 |  | + | - |  | CD122 | 1.01 | 0.84 |  | - | - |
| CD11a | 1.04 | 1.02 |  | - | - |  | CD123 | 1.10 | 0.79 |  | - | - |
| CD11b | 0.92 | 1.06 |  | - | - |  | CD124 | 1.02 | 1.09 |  | - | - |
| CD11c | 0.92 | 1.11 |  | - | - |  | CD126 | 1.03 | 1.02 |  | - | - |
| CD13 | 55.07 | 217.92 |  | + | ++ |  | CD127 | 1.64 | 1.01 |  | - | - |
| CD14 | 0.99 | 1.15 |  | - | - |  | CD132 | 1.03 | 1.56 |  | - | - |
| CD15 | 0.97 | 1.13 |  | - | - |  | CD134 | 0.97 | 1.22 |  | - | - |
| CD16 | 0.95 | 1.08 |  | - | - |  | CD135 | 1.02 | 1.04 |  | - | - |
| CD18 | 0.96 | 1.15 |  | - | - |  | CD137 | 0.96 | 1.15 |  | - | - |
| CD19 | 0.94 | 1.09 |  | - | - |  | CD137L | 0.96 | 1.71 |  | - | - |
| CD20 | 0.97 | 0.90 |  | - | - |  | CD138 | 1.29 | 1.62 |  | - | - |
| CD21 | 0.91 | 1.02 |  | - | - |  | CD140a | 2.79 | 2.44 |  | - | - |
| CD22 | 0.91 | 1.00 |  | - | - |  | CD140b | 20.26 | 2.37 |  | + | - |
| CD23 | 0.92 | 1.06 |  | - | - |  | CD141 | 1.08 | 1.77 |  | - | - |
| CD24 | 1.07 | 1.14 |  | - | - |  | CD144 | 26.88 | 1.25 |  | + | - |
| CD25 | 0.95 | 0.93 |  | - | - |  | CD146 | 17.08 | 14.29 |  | + | + |
| CD26 | 1.85 | 1.81 |  | - | - |  | CD150 | 20.86 | 1.05 |  | + | - |
| CD27 | 1.00 | 1.05 |  | - | - |  | CD152 | 1.07 | 1.01 |  | - | - |
| CD28 | 0.97 | 1.12 |  | - | - |  | CD154 | 0.97 | 44.83 |  | - | + |
| CD29 | 42.31 | 268.79 |  | + | ++ |  | CD158a | 0.99 | 1.05 |  | - | - |
| CD30 | 0.96 | 1.11 |  | - | - |  | CD158b | 1.00 | 1.02 |  | - | - |
| CD31 | 0.88 | 1.13 |  | - | - |  | CD161 | 0.99 | 1.11 |  | - | - |
| CD32 | 0.94 | 1.18 |  | - | - |  | CD162 | 1.09 | 1.24 |  | - | - |
| CD33 | 0.89 | 1.24 |  | - | - |  | CD163 | 1.06 | 0.88 |  | - | - |
| CD34 | 0.98 | 1.13 |  | - | - |  | CD164 | 12.23 | 29.09 |  | + | + |
| CD35 | 0.91 | 1.06 |  | - | - |  | CD165 | 6.66 | 6.63 |  | - | - |
| CD36 | 0.97 | 1.27 |  | - | - |  | CD166 | 27.19 | 123.63 |  | + | ++ |
| CD38 | 0.98 | 1.22 |  | - | - |  | CD172b | 1.13 | 0.77 |  | - | - |
| CD39 | 0.98 | 1.25 |  | - | - |  | CD178 | 1.01 | 0.94 |  | - | - |
| CD40 | 1.05 | 1.22 |  | - | - |  | CD180 | 0.97 | 1.05 |  | - | - |
| CD41 | 1.05 | 1.12 |  | - | - |  | CD181 | 1.06 | 1.73 |  | - | - |
| CD42b | 0.93 | 1.17 |  | - | - |  | CD183 | 1.02 | 1.96 |  | - | - |
| CD43 | 0.92 | 1.53 |  | - | - |  | CD184 | 1.01 | 1.81 |  | - | - |
| CD44 | 454.30 | 536.28 |  | ++ | ++ |  | CD193 | 1.28 | 3.53 |  | - | - |
| CD45 | 1.19 | 1.00 |  | - | - |  | CD195 | 0.99 | 1.06 |  | - | - |
| CD45RA | 0.99 | 0.96 |  | - | - |  | CD196 | 1.00 | 2.22 |  | - | - |
| CD45RB | 0.95 | 0.82 |  | - | - |  | CD197 | 0.96 | 1.05 |  | - | - |
| CD45RO | 0.99 | 0.97 |  | - | - |  | CD200 | 1.00 | 1.85 |  | - | - |
| CD46 | 29.60 | 2.47 |  | + | - |  | CD201 | 3.51 | 5.55 |  | - | - |

|  |  |  |  |  |  |  |  |  |  |  |
| --- | --- | --- | --- | --- | --- | --- | --- | --- | --- | --- |
| CD47 | 30.72 | 32.72 | + | + |  | CD205 | 0.98 | 1.17 | - | - |
| CD48 | 0.97 | 1.29 | - | - |  | CD206 | 0.98 | 0.79 | - | - |
| CD49a | 27.69 | 3.58 | + | - |  | CD209 | 1.09 | 0.78 | - | - |
| CD49c | 71.64 | 75.68 | + | + |  | CD210 | 1.04 | 1.00 | - | - |
| CD49d | 27.06 | 5.66 | + | - |  | CD220 | 1.10 | 1.21 | - | - |
| CD49e | 67.45 | 91.58 | + | + |  | CD221 | 3.94 | 1.98 | - | - |
| CD49f | 37.65 | 35.43 | + | + |  | CD226 | 1.03 | 1.11 | - | - |
| CD50 | 0.99 | 0.99 | - | - |  | CD229 | 1.78 | 0.84 | - | - |
| CD51 | 16.64 | 34.82 | + | + |  | CD231 | 1.04 | 1.11 | - | - |
| CD53 | 0.91 | 1.26 | - | - |  | CD235a | 1.06 | 1.39 | - | - |
| CD54 | 9.41 | 9.68 | - | - |  | CD243 | 1.07 | 1.49 | - | - |
| CD55 | 77.31 | 12.99 | + | + |  | CD244 | 0.96 | 1.08 | - | - |
| CD56 | 2.73 | 1.41 | - | - |  | CD255 | 1.03 | 2.08 | - | - |
| CD57 | 1.10 | 1.01 | - | - |  | CD267 | 0.99 | 1.02 | - | - |
| CD58 | 15.91 | 15.72 | + | + |  | CD268 | 1.06 | 0.99 | - | - |
| CD59 | 122.19 | 1.75 | ++ | - |  | CD271 | 2.28 | 1.05 | - | - |
| CD61 | 17.67 | 1.00 | + | - |  | CD273 | 18.45 | 9.22 | + | - |
| CD62E | 0.92 | 1.08 | - | - |  | CD274 | 14.77 | 3.98 | + | - |
| CD62L | 0.95 | 1.16 | - | - |  | CD275 | 1.03 | 1.09 | - | - |
| CD62P | 0.90 | 55.77 | - | + |  | CD278 | 1.05 | 0.94 | - | - |
| CD63 | 24.50 | 352.06 | + | ++ |  | CD279 | 0.98 | 1.13 | - | - |
| CD64 | 0.90 | 1.13 | - | - |  | CD282 | 0.95 | 0.95 | - | - |
| CD66acde | 0.94 | 1.13 | - | - |  | CD294 | 1.02 | 0.97 | - | - |
| CD66b | 1.02 | 1.12 | - | - |  | CD314 | 0.99 | 1.21 | - | - |
| CD69 | 0.99 | 1.02 | - | - |  | CD326 | 1.12 | 0.92 | - | - |
| CD70 | 1.01 | 8.76 | - | - |  | CDw328 | 0.96 | 0.98 | - | - |
| CD71 | 13.24 | 26.78 | + | + |  | CD335 | 1.01 | 1.05 | - | - |
| CD73 | 95.12 | 75.35 | + | + |  | CD336 | 0.94 | 1.25 | - | - |
| CD74 | 1.05 | 1.29 | - | - |  | CD337 | 0.96 | 1.12 | - | - |
| CD79b | 1.01 | 1.26 | - | - |  | CD338 | 1.19 | 1.09 | - | - |
| CD80 | 1.43 | 1.16 | - | - |  | CD340 | 8.48 | 2.84 | - | - |
| CD81 | 153.86 | 93.47 | ++ | + |  | abTCR | 1.07 |  | - | - |
| CD83 | 0.94 | 1.49 | - | - |  | b2 micro<br>globulin | 45.89 | 199.48 | + | ++ |
| CD84 | 1.04 | 1.22 | - | - |  | BLTR | 0.98 | 1.26 | - | - |
| CD85 | 1.04 | 1.13 | - | - |  | EGF_R | 34.74 | 16.83 | + | + |
| CD86 | 0.96 | 0.95 | - | - |  | HLA_A2 | 1.03 | 1.09 | - | - |
| CD87 | 1.05 | 1.28 | - | - |  | HLA_ABC | 99.57 | 13.81 | + | + |
| CD88 | 0.99 | 0.74 | - | - |  | HLA_DQ | 3.24 | 1.04 | - | - |
| CD89 | 0.93 | 1.13 | - | - |  | HLA_DR | 1.02 | 0.92 | - | - |
| CD90 | 25.98 | 61.91 | + | + |  | Integrin b7 | 1.00 | 0.90 | - | - |
| CD93 | 0.98 | 0.96 | - | - |  | MIC A_B | 2.13 | 2.41 | - | - |
| CD94 | 1.01 | 0.64 | - | - |  | SSEA_1 | 0.95 | 0.95 | - | - |
| CD95 | 25.84 | 17.57 | + | + |  | SSEA_3 | 0.99 | 0.98 | - | - |
| CD97 | 6.09 | 7.95 | - | - |  | SSEA_4 | 3.38 | 7.30 | - | - |
| CD99 | 10.94 | 203.49 | + | ++ |  | TRA_1_60 | 0.91 | 1.10 | - | - |
| CD100 | 1.04 | 1.07 | - | - |  | TRA_1_81 | 1.07 | 1.04 | - | - |
| CD102 | 1.11 | 0.93 | - | - |  | Vb23 | 1.08 | 1.12 | - | - |
| CD103 | 0.93 | 0.77 | - | - |  | Vb8 | 1.01 | 1.05 | - | - |

Data are shown as x-fold expression (MFI) compared to isotype control: +++: >1,000 fold; ++: >100 fold, +: >10 fold, and -: negative vs. isotype control.

Figure 6 illustrates further in vitro characterization of PLX-PAD with the components of PLX-PAD effect on muscle cell proliferation (Figure 6A) and PLX-PAD secretion of factors involved in muscle cell proliferation and migration (Figure 6B).

**Fig. 6.**
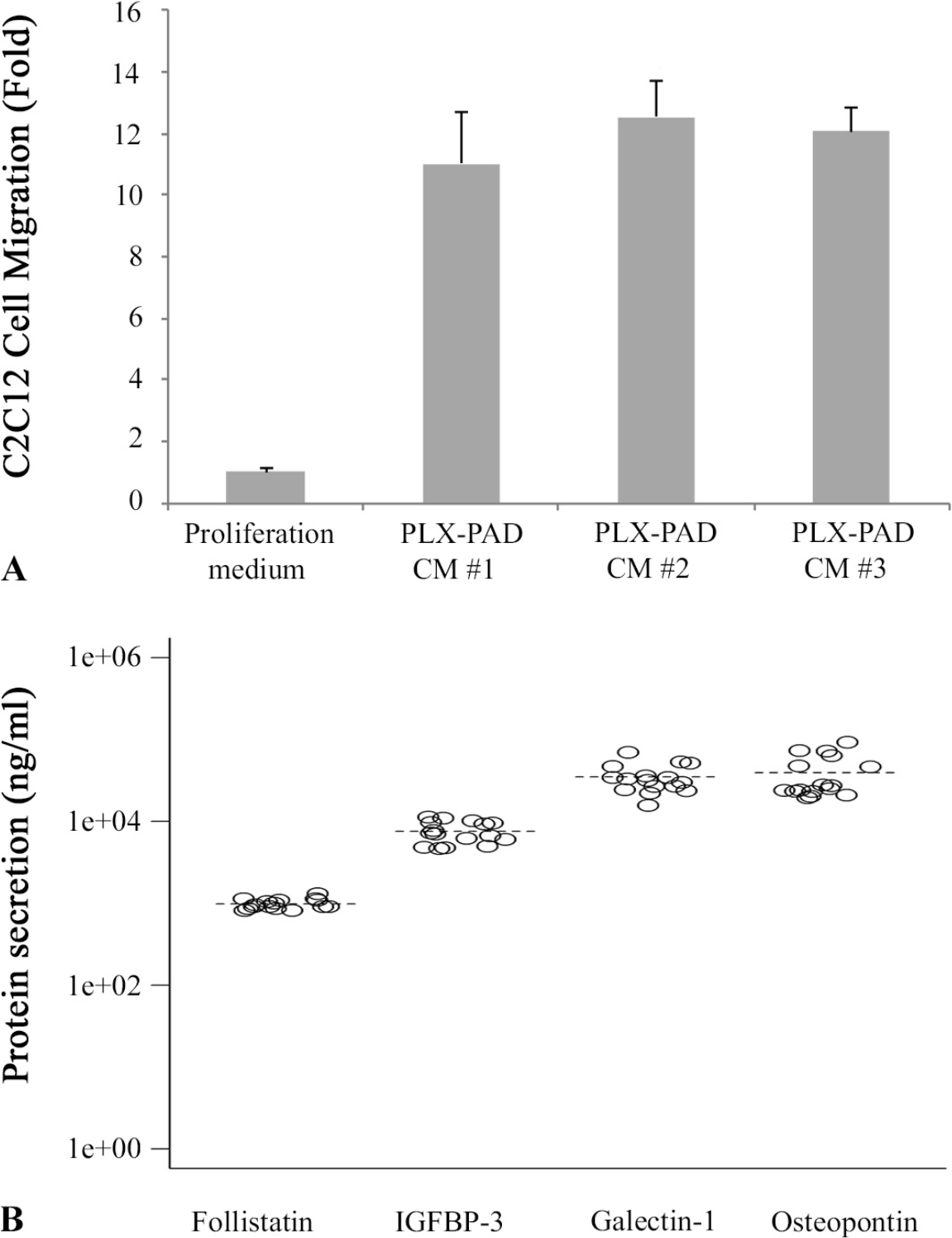
In vitro characterization of PLX-PAD cells. **(A)**Migration of myoblasts (C2C12) incubated with conditioned medium of PLX-PAD cells. CM#1, CM#2 and CM#3 are conditioned media from three batches of PLX-PAD. **(B)** Secretion of Follistatin, IGFBP-3, Galectin-1 and Osteopontin by PLX-PAD in culture.

PLX-PAD cells secrete proteins that are known to be involved in satellite cell activation, proliferation, and migration. Galectin-1, secreted at high levels by PLX-PAD in vitro, is known to be involved in myoblast growth and fusion after muscle injury (*33*), as well as in angiogenesis-related processes (*34, 35*). Osteopontin was shown to be involved in both myogenic and inflammatory processes in early muscle regeneration (*36, 37*). Follistatin is a known regulator of muscle growth and an antagonist of myostatin (which inhibits muscle growth) (*38*). IGFBP-3 belongs to a family of IGF binding proteins that enhance the half-life of IGF, and was specifically shown to support myoblast differentiation and to correlate with increased muscle strength. (*39, 40*).

T cells represent the acquired arm of the immune system, and their activation and proliferation is an important part of the inflammatory process. The effect of PLX-PAD on cell proliferation was assessed in vitro by co-culturing PLX-PAD cells with peripheral blood mononuclear cells (PBMC) stimulated with phytohemagglutinin (PHA), representing a nonspecific T cell mitogen. The results revealed a significant dose dependent decrease in PBMC proliferation (p < 0.001), as shown in Figure 6C.

PLX-PAD were aseptically filled in cryogenic bags at a concentration of 10-20x10^6^ PLX cells/mL in a mixture containing 10% dimethyl sulfoxide (DMSO), 5% human albumin and plasmalyte and stored in gas phase liquid nitrogen at a temperature lower than −150ºC. The required amount of PLX-PAD (1 bag) was thawed in a heated water bath (37°C) immediately prior injection.

### Statistical analysis

Since this was a pilot phase I/IIa trial no formal sample size calculation was performed. We used a modified intention-to-treat (mITT) set including all treated participants. All statistical analyses were performed using SAS (Version 9.2; Cary, North Carolina, USA). We analyzed the biomechanical, macrostructural efficacy endpoints and immunological and hematological parameter changes from baseline (day 0) by applying a mixed model for repeated measures. We analyzed changes in the micro-structural level from baseline until week 12 based on biopsy data using an ANCOVA model. The statistical tests were two-tailed, and we adopted a statistical significance level of P≤0.05.

### Study approval

The study was approved by the German regulatory authorities (Paul Ehrlich Institute, Vorlage- Nr.: 1552101) and the local institutional review board (Landesamt für Gesundheit und Soziales, 12/0045). Written informed consent was collected from all patients, and the study was registered under ClinicalTrials.gov with the identifier NCT01525667 and the European Clinical Trials Database Eudra-CT (2011-003934-16).

## Author Contributions

TW and CP were principal investigators for the study, and designed and managed the study with input from PvR, ANA, BP, MP, SG, ELH, RO, PR, GND and HDV. TW, CP, PvR, ANA, HP, BP, MP, LP, GSD, CM, CC, MS, PR and HDV did the study and collected and/or interpreted data. EE performed a profound statistical reevaluation of the data. TW, GND and HDV drafted the first and subsequent versions of this manuscript with input and key revisions by all authors, who reviewed and approved the final submission.

## Acknowledgements

We thank Anne Zergiebel for her excellent coordination work; Gabriela Korus for her great assistance in histology; Shaul Kadosh for the statistical analysis; Cordula Giesler, Annett Sefrin, Maik Stein and Anke Jurisch for organizing and performing the biomarker tests; and Mark Tastan for assistance with the functional assessments.

## References

1. L. A. Whiteside, Surgical technique: Gluteus maximus and tensor fascia lata transfer for primary deficiency of the abductors of the hip. Clin Orthop Relat Res 472, 645–653 (2014).

2. M. Drexler et al., Acetabular Cup Revision Combined With Tensor Facia Lata Reconstruction for Management of Massive Abductor Avulsion After Failed Total Hip Arthroplasty. J Arthroplasty, (2013).

3. G. M. Alberton, W. A. High, B. F. Morrey, Dislocation after revision total hip arthroplasty: an analysis of risk factors and treatment options. The Journal of bone and joint surgery. American volume 84-A, 1788–1792 (2002).

4. L. A. Whiteside, T. Nayfeh, B. J. Katerberg, Gluteus maximus flap transfer for greater trochanter reconstruction in revision THA. Clin Orthop Relat Res 453, 203–210 (2006).

5. P. von Roth et al., Significant muscle damage after multiple revision total hip replacements through the direct lateral approach. The bone & joint journal 96-B, 1618–1622 (2014).

6. T. Winkler et al., Dose-Response Relationship of Mesenchymal Stem Cell Transplantation and Functional Regeneration after Severe Skeletal Muscle Injury in Rats. Tissue Eng Part A, (2008).

7. T. Winkler et al., Immediate and delayed transplantation of mesenchymal stem cells improve muscle force after skeletal muscle injury in rats. J Tissue Eng Regen Med, (2012).

8. E. K. Merritt et al., Repair of traumatic skeletal muscle injury with bone-marrow-derived mesenchymal stem cells seeded on extracellular matrix. Tissue Eng Part A 16, 2871–2881 (2010).

9. Y. Nakamura et al., Mesenchymal-stem-cell-derived exosomes accelerate skeletal muscle regeneration. FEBS letters 589, 1257–1265 (2015).

10. A. B. Aurora, E. N. Olson, Immune modulation of stem cells and regeneration. Cell stem cell 15, 14–25 (2014).

11. M. Dominici et al., Minimal criteria for defining multipotent mesenchymal stromal cells. The International Society for Cellular Therapy position statement. Cytotherapy 8, 315–317 (2006).

12. S. Barlow et al., Comparison of human placenta- and bone marrow-derived multipotent mesenchymal stem cells. Stem Cells Dev 17, 1095–1107 (2008).

13. R. Roy et al., Cardioprotection by placenta-derived stromal cells in a murine myocardial infarction model. The Journal of surgical research 185, 70–83 (2013).

14. W. R. Prather et al., The role of placental-derived adherent stromal cell (PLX-PAD) in the treatment of critical limb ischemia. Cytotherapy 11, 427–434 (2009).

15. E. Zahavi-Goldstein et al., Placenta-derived PLX-PAD mesenchymal-like stromal cells are efficacious in rescuing blood flow in hind limb ischemia mouse model by a dose- and site-dependent mechanism of action. Cytotherapy 19, 1438–1446 (2017).

16. C. Consentius et al., Mesenchymal Stromal Cells Prevent Allostimulation In Vivo and Control Checkpoints of Th1 Priming: Migration of Human DC to Lymph Nodes and NK Cell Activation. Stem Cells 33, 3087–3099 (2015).

17. W. H. Harris, Traumatic arthritis of the hip after dislocation and acetabular fractures: treatment by mold arthroplasty. An end-result study using a new method of result evaluation. The Journal of bone and joint surgery. American volume 51, 737–755 (1969).

18. J. E. S. Ware K.K.; Kolinski, M.; Gandeck, B, SF-36 Health survey manual and interpretation guide. J. E. Ware, Ed., The Health Institute, New England Medical Centre, Boston, MA (1993).

19. J. Saini, J. S. McPhee, S. Al-Dabbagh, C. E. Stewart, N. Al-Shanti, Regenerative function of immune system: Modulation of muscle stem cells. Ageing Res Rev 27, 67–76 (2016).

20. J. Farup, L. Madaro, P. L. Puri, U. R. Mikkelsen, Interactions between muscle stem cells, mesenchymal-derived cells and immune cells in muscle homeostasis, regeneration and disease. Cell Death Dis 6, e1830 (2015).

21. C. Woiciechowsky et al., Sympathetic activation triggers systemic interleukin-10 release in immunodepression induced by brain injury. Nature medicine 4, 808–813 (1998).

22. E. Li Causi et al., Vaccination Expands Antigen-Specific CD4+ Memory T Cells and Mobilizes Bystander Central Memory T Cells. PloS one 10, e0136717 (2015).

23. J. P. Desborough, The stress response to trauma and surgery. British journal of anaesthesia 85, 109–117 (2000).

24. J. P. Farthing, E. P. Zehr, Restoring symmetry: clinical applications of cross-education. Exerc Sport Sci Rev 42, 70–75 (2014).

25. T. Partridge, The current status of myoblast transfer. Neurological sciences: official journal of the Italian Neurological Society and of the Italian Society of Clinical Neurophysiology 21, S939–942 (2000).

26. D. Montarras et al., Direct isolation of satellite cells for skeletal muscle regeneration. Science 309, 2064–2067 (2005).

27. P. Hong et al., HEXIM1 controls satellite cell expansion after injury to regulate skeletal muscle regeneration. J Clin Invest 122, 3873–3887 (2012).

28. C. H. Lee et al., Harnessing endogenous stem/progenitor cells for tendon regeneration. J Clin Invest 125, 2690–2701 (2015).

29. Y. Ramot, M. Meiron, A. Toren, M. Steiner, A. Nyska, Safety and biodistribution profile of placental-derived mesenchymal stromal cells (PLX-PAD) following intramuscular delivery. Toxicologic pathology 37, 606–616 (2009).

30. D. Sugimoto, C. G. Mattacola, D. R. Mullineaux, T. G. Palmer, T. E. Hewett, Comparison of isokinetic hip abduction and adduction peak torques and ratio between sexes. Clin J Sport Med 24, 422–428 (2014).

31. J. A. Mutchler, J. T. Weinhandl, M. C. Hoch, B. L. Van Lunen, Reliability and fatigue characteristics of a standing hip isometric endurance protocol. J Electromyogr Kinesiol 25, 667–674 (2015).

32. F. Engelken et al., Assessment of fatty degeneration of the gluteal muscles in patients with THA using MRI: reliability and accuracy of the Goutallier and quartile classification systems. The Journal of arthroplasty 29, 149–153 (2014).

33. K. Kami, E. Senba, Galectin-1 is a novel factor that regulates myotube growth in regenerating skeletal muscles. Current drug targets 6, 395–405 (2005).

34. D. O. Croci et al., Disrupting galectin-1 interactions with N-glycans suppresses hypoxia-driven angiogenesis and tumorigenesis in Kaposi’s sarcoma. The Journal of experimental medicine 209, 1985–2000 (2012).

35. D. O. Croci et al., Glycosylation-dependent lectin-receptor interactions preserve angiogenesis in anti-VEGF refractory tumors. Cell 156, 744–758 (2014).

36. A. Hirata et al., Expression profiling of cytokines and related genes in regenerating skeletal muscle after cardiotoxin injection: a role for osteopontin. The American journal of pathology 163, 203–215 (2003).

37. K. Uaesoontrachoon et al., Osteopontin and skeletal muscle myoblasts: association with muscle regeneration and regulation of myoblast function in vitro. The international journal of biochemistry & cell biology 40, 2303–2314 (2008).

38. S. J. Lee, A. C. McPherron, Regulation of myostatin activity and muscle growth. Proceedings of the National Academy of Sciences of the United States of America 98, 9306–9311 (2001).

39. E. J. Foulstone, P. B. Savage, A. L. Crown, J. M. Holly, C. E. Stewart, Role of insulin-like growth factor binding protein-3 (IGFBP-3) in the differentiation of primary human adult skeletal myoblasts. Journal of cellular physiology 195, 70–79 (2003).

40. D. G. Taekema et al., Circulating levels of IGF1 are associated with muscle strength in middle-aged- and oldest-old women. European journal of endocrinology / European Federation of Endocrine Societies 164, 189–196 (2011).

